# Optimal choice of parameters in functional connectome-based predictive modelling might be biased by motion: comment on Dadi et al

**DOI:** 10.1101/710731

**Authors:** Tamas Spisak, Balint Kincses, Ulrike Bingel

## Abstract

In a recent study, Dadi and colleagues make recommendations on optimal parameters for functional connectome-based predictive models. While the authors acknowledge that “optimal choices of parameters will differ on datasets with very different properties”, some questions regarding the universality of the recommended “default values” remain unanswered.

Namely, as already briefly discussed by Dadi et al., the datasets used in the target study might not be representative regarding the sparsity of the (hidden) ground truth (i.e. the number of non-informative connections), which might affect the performance of L_1_- and L_2_-regularization approaches and feature selection.

Here we exemplify that, at least in one of the investigated datasets systematic motion artefacts might bias the discriminative signal towards “non-sparsity”, which might lead to underestimating the performance of L_1_-regularized models and feature selection.

We conclude that the expected sparsity of the discriminative signal should be carefully considered when planning predictive modelling workflows and the neuroscientific validity of predictive models should be investigated to account for non-neural confounds.

## Introduction

In a recent study, Dadi and colleagues^1^ systematically compare different analysis pipelines for predictive modelling based on the resting-state functional connectome. They obtain imaging and phenotypic data from six different datasets with a total of 2000 individuals and make recommendations regarding optimal parameter choices for predictive modelling.

Dadi and colleagues recommend, among others, the use of non-sparse (L_2_-regularized) modelling approaches, albeit they also briefly highlight that “the lack of success of sparse approaches suggests that the discriminant signal is distributed across the functional connectome for the tasks we study”. Indeed, it seems plausible that the representation of various conditions of interest can widely differ in the functional connectome; some might be driven by a limited (“sparse”) set of “discriminative” connections, others by distributed, “non-sparse” patterns.

Importantly, sparsity and other important features of the discriminative ground truth can be confounded by signal of non-neural origin. E.g. motion artefacts, a well-known source of confounds in functional connectivity analysis^2^, are known to introduce complex spatio-temporal patterns in the resting-state data and can systematically differ between groups or conditions of interest. In fact, we have previously reported^3^ such systematic motion artefacts in the Autism Brain Imaging Data Exchange (ABIDE) dataset^4^, which was also investigated in the target study.

Here, we hypothesize that certain methods in predictive modelling might be prone to learn the typical “non-sparse”, global effects of motion artifacts and possibly bias the results of the target study.

To test this hypothesis, we used the source code nobly shared by the authors and investigated whether a subject-level summary descriptor of in-scanner head motion is correlated with predicted class labels in the ABIDE dataset. Using this value as a proxy for the “artefact-dependence” of prediction, we contrasted sparse and non-sparse regularization techniques and various types of connectivity estimates.

## Methods

Cross-validated training and test of the *L*_*1*_ and *L*_*2*_-regularised logistic regression models were performed on correlation, partial correlation and tangent values computed on the BASC atlas-based timeseries data from the ABIDE dataset. Timeseries were obtained as published by Dadi et al. (https://github.com/KamalakerDadi/benchmark_rsfMRI_prediction). Analysis was identical to that used for Fig. 7 of the target article, using 100 random, stratified splits (75% training data, 25% test data) to obtain ROC AUC values.

Mean framewise displacement (FD) for each subject was obtained from the phenotypical datasheet of the preprocessed ABIDE database^5^. Difference in group-averaged mean FD between the patient and control groups was computed. Mean FD difference was also computed between the predicted classes in each random stratified fold and contrasted to the mean FD difference based on the diagnosis to estimate the extent to which the prediction captured systematic motion-artefacts. Corresponding source code is shared at https://github.com/spisakt/benchmark_rsfMRI_prediction.

## Results and Discussion

We have found that in the ABIDE subsample used in the target study (N=866), motion (mean FD) significantly differs between the diagnoses (ΔmeanFD=0.03mm, p<0.0001, dashed line on Figure 1), with patients moving more than controls.

**Figure 1.**
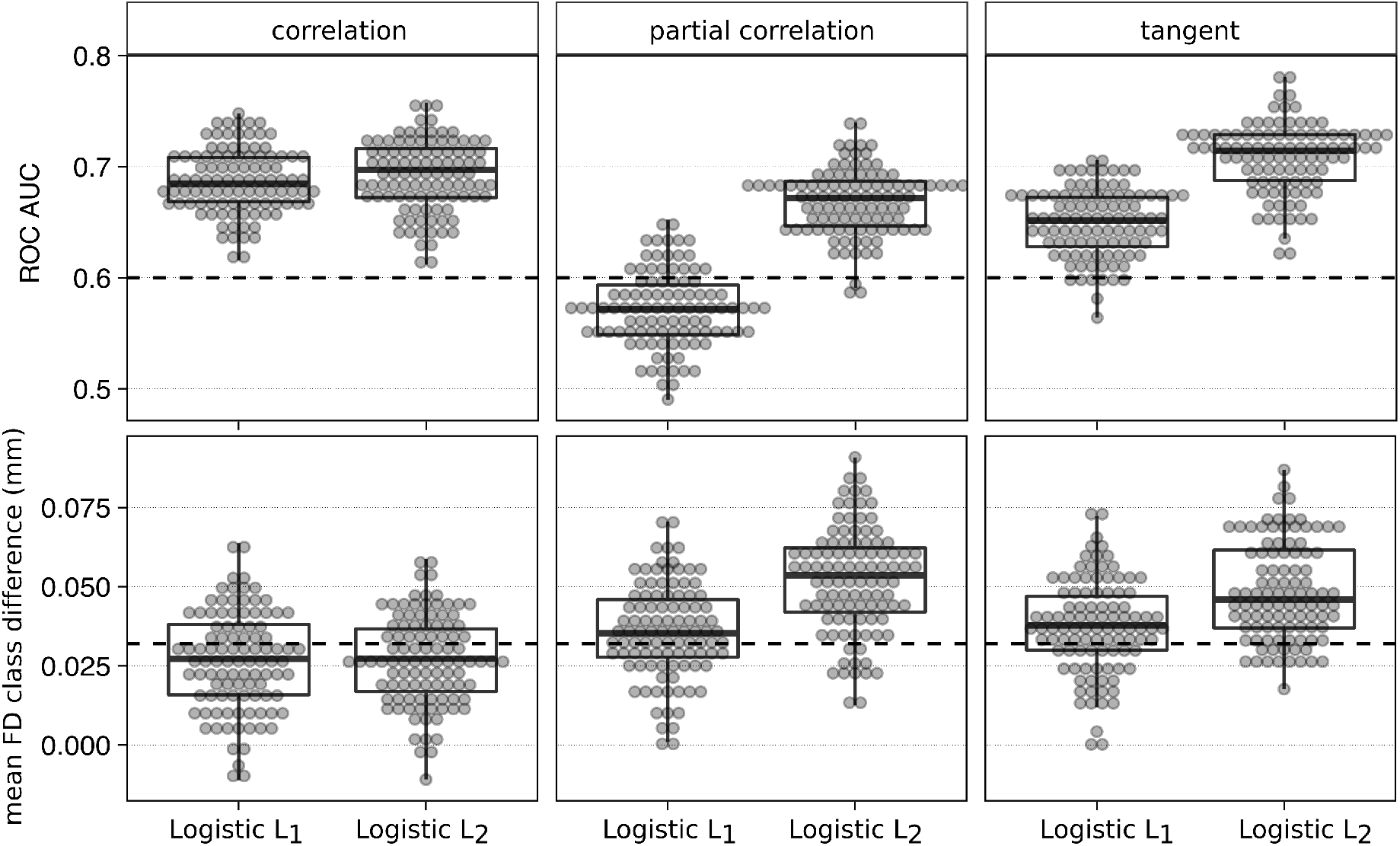
Better performance of certain predictive analysis workflows might be partially driven by globally-distributed motion artefacts in the ABIDE dataset. The upper row depicts the performance of the L_1_- and L_2_-regularused logistic regressions in the ABIDE dataset, with correlation, partial correlation and tangent-space embedding as the measure of functional connectivity, throughout the cross-validation splits. Dashed line denotes the ROC AUC value (0.6) of the classification entirety based on motion (FD). The bottom row shows the difference in mean framewise displacement between the predicted classes (as a proxy for motion artefacts) in the same data, across the same cross-validation splits. Dashed line denotes the difference in mean framewise displacement between the groups based on the actual diagnoses (ΔmeanFD=0.03mm, p<0.0001). For all plots, the same subjects, region-definition (BASC-network), timeseries and cross-validation strategy was used as for the corresponding parts of Figure 7 of the target paper.

With such a difference, individual mean FD, alone, is able to predict the diagnosis with an ROC AUC=0.6 (Figure 1, upper row, dashed line).

While in the investigated sample, L_2_-regularization with tangent-space embedding tended to outperform other pipeline settings (see Figure 7 of the target article^1^ and the upper row of our Figure 1), we have found that this model (L_2_-regularization with tangent-space embedding but also partial correlation) also resulted in strongly motion-correlated predictions. Importantly, the motion difference between the predicted classes was significantly greater than for the true class labels (Figure 1, bottom row), strongly suggesting that the relatively good performance of this models is partially driven by motion-associated connectivity differences.

Although these motion-related differences could be also of neural origin, as we previously demonstrated^3^ in the ABIDE dataset, motion exerts its artifactual effect in the form of complex, distributed (i.e. non-sparse) spatio-temporal artefact-patterns, which systematically differ between the groups of interest.

Therefore, a plausible interpretation of our results is that the gain in the predictive performance with non-sparse regularization approaches might partially be driven by their ability to capture non-sparse, motion-related confounder signal.

Since autism spectrum disorder is known to affect a large part of the functional connectome^3,4^, it is implausible that the gain in prediction performance with L_2_-regularization is exclusively driven by motion artefacts. On the other hand, the discriminative signal in some states or traits of interest might follow a much sparser representation^6–9^. In these cases, L_1_-regularization and/or feature selection might be advantageous not only for known theoretical reasons^10–13^, but might also achieve a better neuroscientific validity than non-sparse approaches, by dropping the spatially distributed but weak effects of motion artefacts.

## Conclusion

We strongly appreciate the authors’ effort to provide methodological guidance for functional connectivity-based predictive modelling. Such efforts are urgently needed to avoid the combinatorial explosion of computation costs and to overcome unwanted proliferation of the corresponding methodology. We believe that their recommendations will significantly foster future research efforts in search for robust and reliable biomarkers.

Here we argue that motion-related confounds might contribute to the discriminative connectivity signal as a non-sparse non-neural component and, accordingly, can be mitigated by the use of sparse modelling approaches or feature selection.

Therefore, we conclude that the expected sparsity of the discriminative features and the potentially better neuroscientific validity of sparse models should be carefully considered when drawing decisions about the analysis strategy in predictive modelling.

## Acknowledgements

We are thankful to Dadi and colleagues for publishing the source code of their analysis. This commentary is fundamentally built on this code and, therefore, demonstrates how transparency and availability of the methodology substantially fosters objective and informed scientific arguments.

B.K is supported by the UNKP-18-3 New National Excellence Program of the Ministry of Human Capacities, Hungary.

